# Protein Structure Refinement via DeepTracer and AlphaFold2

**DOI:** 10.1101/2023.08.16.553616

**Authors:** Jason Chen, Ayisha Zia, Fengbin Wang, Jie Hou, Renzhi Cao, Dong Si

## Abstract

Understanding the structures of proteins has numerous applications, such as vaccine development. It is a slow and labor-intensive task to manually build protein structures from experimental electron density maps, therefore, machine learning approaches have been proposed to automate this process. However, most of the experimental maps are not atomic resolution, so they are insufficient for computer vision-based machine learning methods to precisely determine the protein structure. On the other hand, methods that utilize evolutionary information from protein sequences to predict structures, like AlphaFold2, have recently achieved groundbreaking accuracy but often require manual effort to refine the results. We propose DeepTracer-Refine, an automated method to refine AlphaFold structures by aligning them to DeepTracer’s predicted structure. We tested our method on 39 multi-domain proteins and we improved the average residue coverage from 78.2% to 90.0% and average lDDT score from 0.67 to 0.71. We also compared DeepTracer-Refine against another method, Phenix’s AlphaFold refinement, to demonstrate that our method not only performs better when the initial AlphaFold model is less precise but also exceeds Phenix in run-time performance.

## Introduction

Macromolecular imaging technology like Cryogenic electron microscopy (cryo-EM) has revolutionized the way we study proteins [1]. Proteins perform many important functions within the human body and we study their three-dimensional composition to gain knowledge about their functions [2]. In the past few years, many *de novo* and template-based machine learning methods have been proposed to automate the process of protein structure modeling from cryo-EM density maps [3–9] but they have challenges in producing an accurate structure. For *de novo* methods, meaning that they do not rely on using previously solved structures as templates, they are constrained by the resolution and quality of the density maps. For template-based methods, they are constrained by the availability of solved structures. These limitations could be overcome with better electron microscopes and growth in the availability of solved structures, however, from a computer algorithm perspective, we need to look elsewhere to overcome the limitations of current automated methods.

There are two general approaches when it comes to determining protein structures *in silico*. The first is map-to-model, which places atoms in experimentally obtained cryo-EM electron density maps (Figure 1A); the second is sequence-to-model, which utilizes information from the protein sequence to predict the three-dimensional structure (Figure 1B). We propose DeepTracer-Refine to address the limitations of our map-to-model method, DeepTracer, by incorporating it with a sequence-to-model method, AlphaFold2. In short, AlphaFold structures provide perfect sequence coverage because it uses the true protein sequence as input. However, AlphaFold’s predicted structures are not as precise compared to *de novo* map-to-model methods like DeepTracer. A common strategy to address this problem is by splitting AlphaFold’s structure into compact domains, i.e. well folded parts, and manually docking them into experimental maps to fix the residue locations. DeepTracer-Refine is an automated method that splits AlphaFold’s prediction into compact domains and aligns them to DeepTracer’s prediction instead of an experimental map.

**Figure 1.**
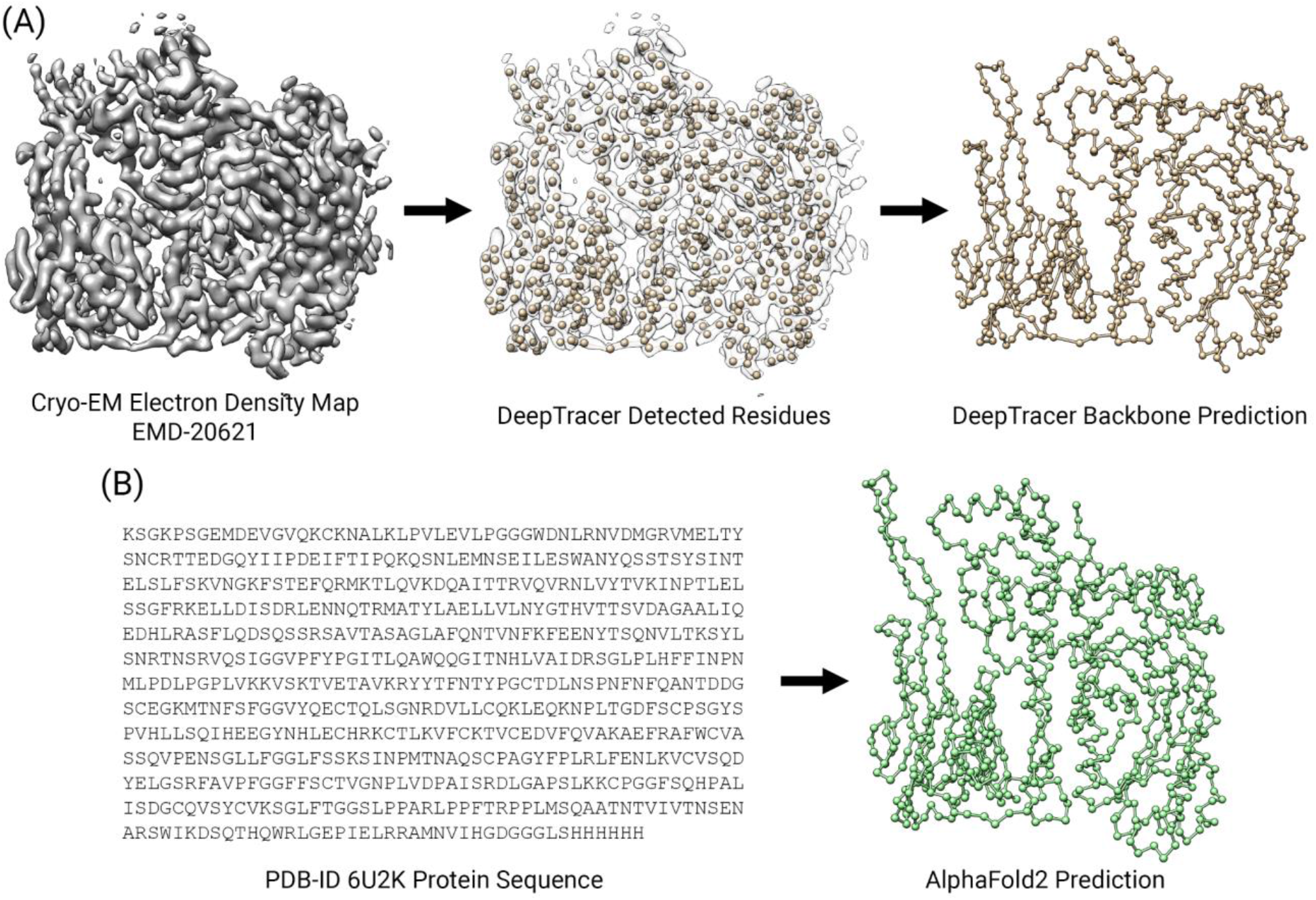
An illustration of the two approaches in protein structure prediction. **(A)** Map-to-model detects the residues from a cryo-EM electron density map and connects them into a 3D structure through backbone tracing. **(B)** Sequence-to-model uses a protein sequence as input to fold the 1D sequence into a 3D structure.

### Protein Structure

Briefly, a protein structure consists of one or multiple chains of amino acids. When two amino acids are bonded together through a peptide bond, a water molecule is released and the remaining components are called residues. Thus, we can consider a residue as the basic unit of a protein structure. A polypeptide chain is a sequence of many amino acids bonded together in a linear fashion to form the three-dimensional shape of a protein structure. There are 20 types of amino acids observed in natural protein sequences and each comprising a backbone and a side-chain portion. The backbone of every amino acid is composed of the same atoms that link together to form the “skeleton” of the structure. However, it is the side chain that differentiates an amino acid’s type.

### Challenges of Map-to-Model Approaches

The main objective of map-to-model methods is to identify the type and location of residues and then connect them into a three-dimensional structure. Map-to-model methods often use machine learning-based computer vision algorithms to detect the residues from experimental maps, however, they use various strategies to connect the residues. In order to predict an accurate structure, the type of residues must be correctly identified and then connected in the proper sequential order. *De novo* methods commonly employ heuristic path-walking and tree-graphs algorithms to connect the residues without requiring a database of template structures [3–6]. Currently, it is a challenge to accurately identify amino acid types because the resolution of cryo-EM is not yet true atomic so machine learning has difficulty classifying the side chains. As a consequence, *de novo* methods typically have to post-process the initial result to improve the sequence prediction, such as DeepTracer.

DeepTracer utilizes a 3D convolutional neural network called the U-Net to identify residues from cryo-EM density maps. It and other non-template-based *de novo* methods are only capable of identifying residues from the region of experimental maps that cover the protein. This means that if the map does not fully cover the protein, then DeepTracer will not be able to predict the full structure. Experimental maps often do not cover the entire structure of the imaged protein, therefore, DeepTracer’s predicted structural coverage is often incomplete. This is also the reason why many manually solved structures contain missing residues. In addition, experimental maps are noisy in nature and they come in many sizes and qualities. DeepTracer has attempted to address these problems by pre-processing the density maps into consistent input, however, It is impossible to train a machine learning model that works on every input without overfitting. On high-resolution and high-quality density maps like EMD-20621 [10], DeepTracer can achieve above 90% residue coverage, however, it is the missing 10% that creates challenges for DeepTracer.

The first challenge is false connections. The process of connecting residues is called backbone tracing and the order in which the residues are connected determines the predicted protein sequence. If the residues are connected in the wrong order, then the sequence would also be out-of-order and thus incorrect. DeepTracer employs a traveling salesman algorithm to trace the backbone, which connects closest residues based on Euclidean coordinates. However, the issue arises when residues are missing and it causes a faulty connection in the backbone. In these cases, the nearest residue in space is often not the next closest sequential residue and we can see this in the example of EMD-20621 (Figure 2B).

**Figure 2.**
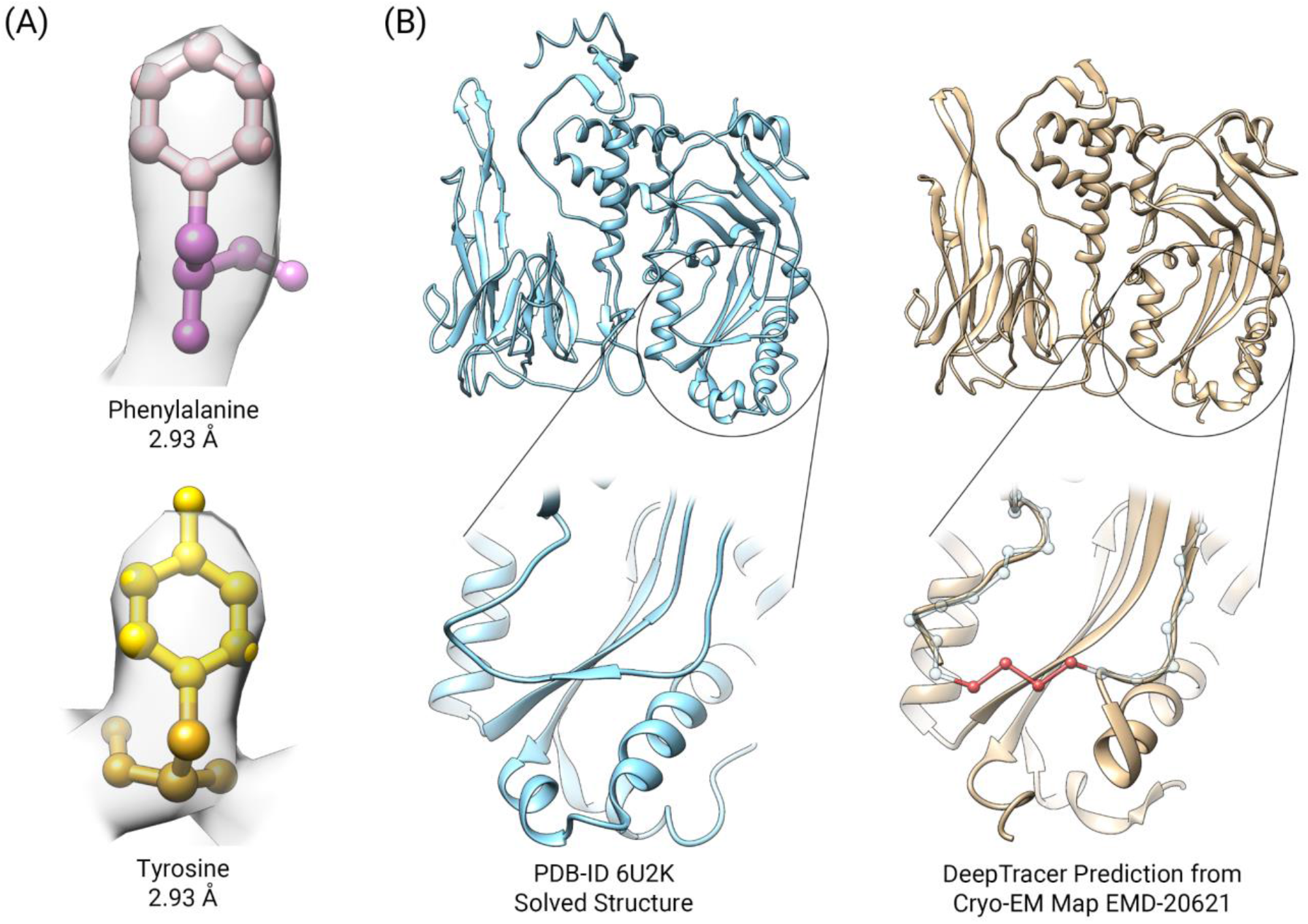
Current challenges of map-to-model methods. **(A)** Cryo-EM maps lack the resolution to provide atomic-level details for accurate side-chain identification even at high resolution. Two residues, tyrosine and phenylalanine, are placed in cryo-EM density map EMD-20621 and the side-chains are indistinguishable from the map. The side-chain atoms are colored in lighter colors respectively. **(B)** Missing residues causing false connections. In the close-up section, DeepTracer missed four residues (overlay highlighted in red) causing the backbone to be traced incorrectly.

**Figure 3.**
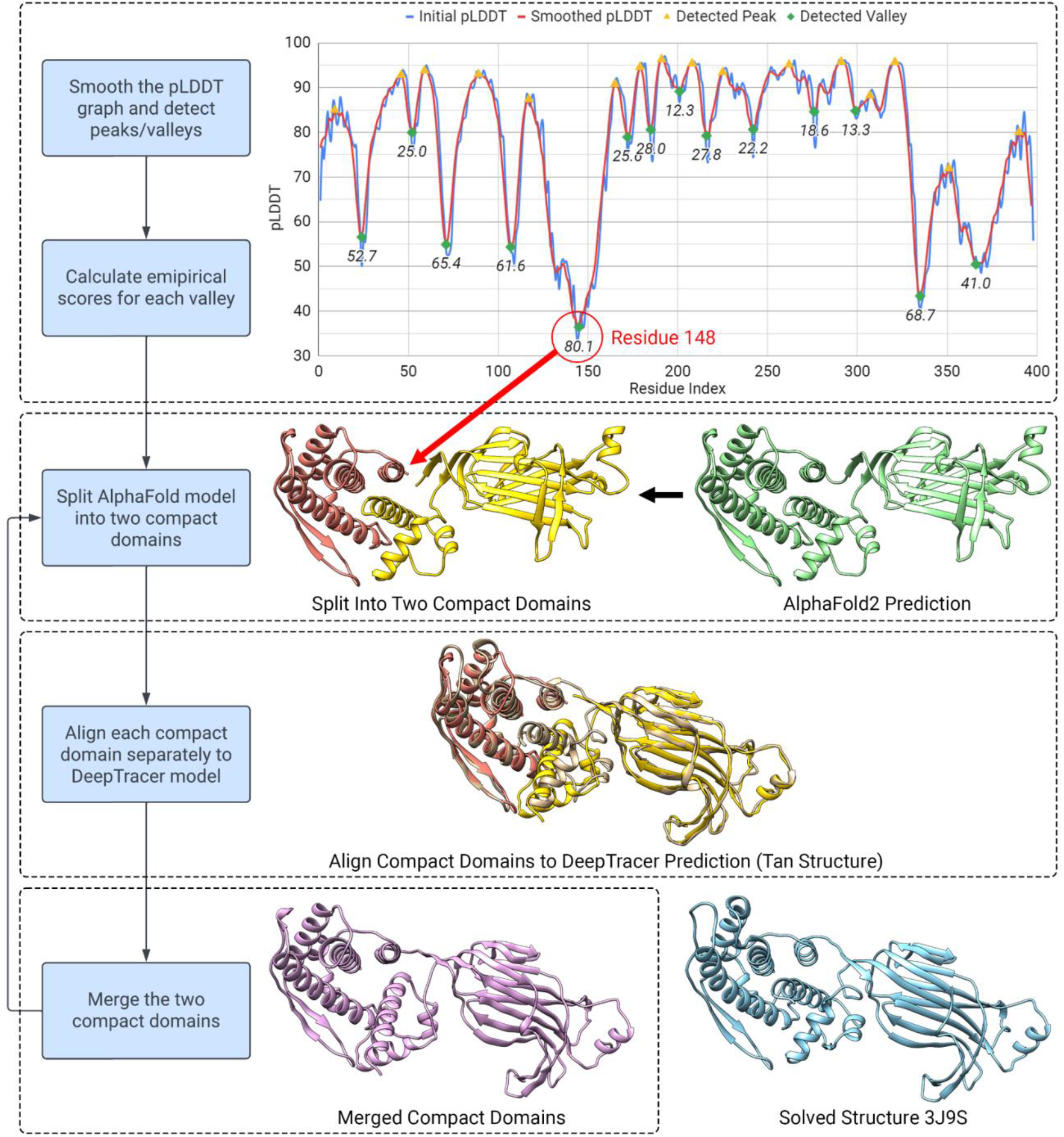
The design of DeepTracer-Refine pipeline. We utilized AlphaFold’s pLDDT score to detect and rank locations to split the structure into compact domains. We process the pLDDT scores first and calculate an empirical score for each possible low-confidence location. We rank the locations based on the empirical scores and a higher score suggests a poorer AlphaFold prediction. In this example of a rotavirus VP6 [27], we split the AlphaFold model at residue 148 and aligned each domain separately to DeepTracer’s prediction. The merged structure is improved as we can see the conformation of the right-side structure resembles more closely to the solved structure.

The second challenge is the predicted sequence accuracy. Although DeepTracer can effectively identify residues from near-atomic cryo-EM density maps, it cannot accurately classify the types due to resolution limits. Current high resolution maps have a resolution around 2 to 4 Å, however, the diameter of an atom is approximately 1 Å so it is insufficient to distinguish individual atoms (Figure 2A). DeepTracer classifies residue types by the shape of its side-chain atom group and there is not enough detail even at high resolution to clearly differentiate the side-chain shapes. Since the sequence is determined by the amino acid types and their sequential order, false connections and incorrect type predictions result in an inaccurately predicted sequence. DeepTracer solves this problem by performing a sequence alignment between the predicted sequence and true sequence. It uses dynamic programming to find the best alignment between the two sequences and then replaces the predicted sequence with the true sequence. The sequence alignment is effective but it is dependent on the quality of the backbone trace, in which the false connections can influence its effectiveness.

### Challenges of Sequence-to-Model Approaches

The main objective of sequence-to-model is to fold protein sequences into a three-dimensional conformation using evolutionary information and energy functions. A protein sequence is a linear arrangement of amino acids like words in a sentence. Every type of amino acid possesses different side-chain charges and they exert forces on each other when bonded together. The 1D sequence is inclined to fold into a stable 3D conformation with the lowest energy landscape based on Anfinsen’s thermodynamic hypothesis [11]. As such, proteins fold into distinctive shapes depending on the combination of amino acids and thus the sequences embed rich information regarding their structure. Methods have been proposed to utilize natural language processing techniques to extract useful information from protein sequences [12–15]. However, deriving the 3D structure through an energy-guided approach presents a large search space [16] so prior sequence-to-model methods were only able to fold small to medium proteins (less than 800 residues) [17, 18].

AlphaFold2, which we will refer to as AlphaFold, is the second version of AlphaFold and it achieved groundbreaking accuracy in the 14th Critical Assessment of Structure Prediction (CASP14) conference [19]. It and other similar sequence-to-model methods [20] have recently adopted a new kind of machine-learning architecture termed the transformer [21]. The transformer has a unique attention mechanism capable of learning contextualization, and therefore, significantly reducing the search space for an energy-based approach [22]. AlphaFold also combined evolutionary information by using multiple sequence alignment to further achieve state-of-the-art accuracy. Since it uses true sequence as the input, it does not suffer from the same residue coverage and sequence prediction issues that map-to-model methods do. The main challenge of AlphaFold is accurately folding the regions between protein domains. In structural biology, a protein domain is a spatially distinct unit that is folded in a compact conformation [23]. AlphaFold models are often less confident at the regions in-between compact domains due to the flexible and intrinsically disordered nature of these regions [18, 24].

### DeepTracer-Refine Pipeline

Map-to-model and sequence-to-model have their own advantages which complement each other’s challenges. So we developed DeepTracer-Refine to take advantage of the perfect sequence of AlphaFold structures and improve the less accurate backbone regions by utilizing DeepTracer’s prediction. DeepTracer-Refine is an automated pipeline that detects optimal locations to split the AlphaFold structure into compact domains and make iteratively alignments to improve the predicted structure. AlphaFold generates a per-residue metric called the predicted Local Distance Difference Test (pLDDT) to estimate the confidence level of its prediction for each residue [19]. It has been shown that pLDDT correlates highly to the accuracy of AlphaFold’s predicted structure [22] so we used it to determine where to split the structure. In a pLDDT graph, low-confidence regions are shaped like valleys and we observed that a steeper and deeper valley is often a better indicator of an optimal splitting spot than the raw pLDDT score.

Since we are iteratively splitting and aligning compact domains, the order and location in which we split the AlphaFold prediction can affect the final result. We introduce movements to the entire structure when we align each compact domain, therefore, we want to minimize movements towards the end. We developed an empirical score based on pLDDT to quantify the poorness of the fold so we can rank them for splitting. We treat the pLDDT as a signal and process it in three steps: (1) smoothing the signal, (2) finding the prominent peaks and valleys, and (3) obtaining a score for each valley. We first process the signal by smoothing it with a 1D filter to remove noise so the peak detection is more reliable. Secondly, we apply Scipy’s `find_peaks` function to detect the peaks (local maximas) and valleys (local minimas). We use a prominence setting of six to ensure we only get the significant local maximas and minimas. A wider peak-to-peak distance indicates a valley is less steep, so we penalize it by subtracting the valley width from 200 (Equation 2). We use the average depth (Equation 1) and the valley width to calculate the final empirical score (Equation 3). A higher score suggests a less accurate fold.
>

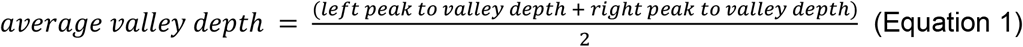

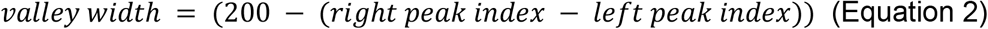

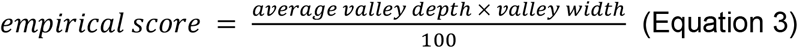

We adopted an iterative alignment strategy, so we are not limited to a fixed amount of alignments. The refinement process is a split-align-merge procedure, which we attempt for every possible detected splitting location. The order in which we split the model is determined by the empirical score from highest to lowest. For each round, we split the model into two compact domains and align them separately to DeepTracer’s structure. We align the larger domain first so the smaller one will have a better chance to align to the correct position. If the aligned domains are not too distant to be connected, then we merge them together and calculate the residue coverage. If the residue coverage is increased by more than a threshold of 1%, then we consider it an improvement and we keep the refined structure; otherwise, we keep the previous structure. We also avoid splitting if the segment length of a domain is less than 40 residues. The segments cannot be too short because false alignments are more likely to occur. The result from the last round of the iterative split-align-merge process is the final DeepTracer-Refine output.

Since the compact domains could be aligned to any parts of DeepTracer’s prediction, we employ two strategies to prevent false alignments. The first issue is overlapping alignments to which the compact domains are aligned to the same location on DeepTracer’s structure. Once we align the first domain, we remove the matching residues from DeepTracer’s structure to prevent the second domain from aligning to the same location. The second issue is distant alignment in which two domains are aligned so far apart that they cannot be physically connected. The intrinsically disordered regions are often loop sections so we check if there are enough residues to span the distance in-between the last non-loop residues from the ends of the compact domains. We count the number of residues of the loop section, multiply it by 3 Å, which is slightly less than the average distance between residues, and compare it to the distance between the last non-loop residues. According to our tests, both strategies decreased the number of false alignments when using heuristic alignment algorithms.

A flexible structure alignment algorithm seems like an appropriate solution for fixing the inaccurately folded regions. FATCAT [24] is a flexible alignment method but we discovered that DeepTracer’s false connection and imperfect sequence prediction often led to many incorrectly transformed fragments. Since we are already splitting the AlphaFold model into shorter segments, we decided to use rigid body alignment methods, which includes PyMOL cealign [25], PyMOL align, and Chimera MatchMaker [26]. We run all three algorithms on each compact domain and select the one with the highest residues coverage after merging. The AlphaFold’s residue locations are updated when each domain is aligned to DeepTracer’s prediction.

### Experiment Details

We used a local version of ColabFold [28] to generate AlphaFold predictions. ColabFold with the default settings produces five models and we used the rank one model for our testing. We ran DeepTracer-Refine and the PHENIX software suite (version 1.20.1-4487) on a Windows machine with an Intel Core i7-4790 8-core CPU, 16GB of ram, and an Nvidia GeForce GTX 780 Ti graphics card. All of the protein structures and cryo-EM density maps are visualized through UCSF Chimera [29].

### Data Set

We retrieved the cryo-EM density maps, solved structures, and protein sequences from the RCSB protein data bank using PDB-IDs [30]. Since we are aligning AlphaFold models to DeepTracer structures, the target must first contain a corresponding cryo-EM density map. DeepTracer is optimized to work on near-atomic resolution maps so we filtered the search to cryo-EM with a resolution of 3.5 Å or higher. We then limited the number of residues per target from 200 to 1000 to ensure the protein is not too small and the ColabFold run time is within a reasonable range. We only tested on monomeric proteins, i.e., single-chain proteins, otherwise compact domains can be aligned to more than one location in a symmetrical multimer with duplicate copies of chain(s). We conducted a query based on these parameters via RCSB advanced search and it returned 59 targets. After manually removing the multimers and duplicates, we were left with a total of 39 targets.

### Evaluation Metric

In order to evaluate DeepTracer-Refine, we used the solved structure as ground truth for our comparisons and utilized two metrics to evaluate the results. The first is overall structural coverage in the form of residue matching percentage (Equation 4). We consider a predicted residue and a solved residue matching if they are within a 3 Å distance. Since residue matching relies on superpositioning, we used Chimera MatchMaker to align the AlphaFold prediction to the solved structure to calculate the initial residue coverage. We assume that if the matching percentage increased after DeepTracer-Refine, then we have improved the initial AlphaFold structure. However, residue coverage does not convey structural similarity so we use the local Distance Difference Test (lDDT) score as the second metric.

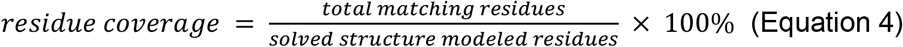

Since we are only comparing the backbone geometry, we use Cα-lDDT which only considers the Cα atom for each residue. The structural similarity between a solved and predicted residue is quantified by the fraction of preserved distances with regards to their respective neighboring residues [31]. The size of the neighborhood is determined by the inclusion radius. The default is 15 Å because it allows lDDT to ignore movements of compact domains, meaning that two structure can be viewed as similar even if the corresponding domains are not in the same relative location. However, we are attempting to address the imprecisely folded flexible regions so we need to set a large inclusion radius to ignore domain movements. We chose an inclusion radius of 200 Å to include all pairs of residues in our evaluation. A distance is considered preserved if the difference between the same pair of residues from the predicted and solved structure is within a threshold. The final lDDT score is an average of the results from four thresholds, 0.5 Å, 1 Å, 2 Å, and 4 Å.

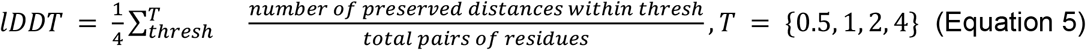

## Results

We first compared the results of DeepTracer-Refine to AlphaFold’s initial predictions to evaluate the effectiveness of our method. DeepTracer-Refine increased the overall average residue coverage from 77.8% to 90.0% and average lDDT from 0.667 to 0.707. Out of the 39 targets, it improved 27 of the structures according to the increase in both residue coverage and lDDT. There were also eight unimproved and four worsened structures. The average residue coverage and average lDDT for the unaltered and unimproved structures were 95.84% and 0.799. We argue that they were already adequate for manual refinement and thus do not require DeepTracer-Refine. Although the residue coverage increased for all but five models, there were a total of seven that decreased in lDDT scores and four out of the seven with visible negative structural changes. For the full list of the result, please refer to the supplementary section.

One of the most improved structures was a human norovirus GII.2 Snow Mountain Virus strain VLP asymmetric unit (Figure 4A) [32]. We saw improvements in both the residue coverage and lDDT score, with an increase of 32.7% in residue coverage and 0.321 in lDDT. Initially, the top-left domain of the AlphaFold model from residues 1 to 218 was incorrectly folded. DeepTracer-Refine split the AlphaFold model at residue 218 and aligned each half correctly. The original DeepTracer prediction was very accurate in this case and it achieved 97.5% residue coverage. The DeepTracer backbone did not contain any false connections so it effectively aligned the sequence and improved the predicted sequence accuracy to 93.5%. In another case, a human OCT3 in lipid nanodisc [33], the sequence prediction of DeepTracer was not as accurate as the previous example. DeepTracer achieved 88.4% residue coverage but only attained 59.8% sequence accuracy even after sequence alignment. This was mainly due to false connections which reduced the effectiveness of the sequence alignment post-processing step. In Figure 4B, the red encircled region represents residues 330 to 554, and there is a noticeable difference in the conformation when compared to the solved structure 7ZH0. DeepTracer-Refine improved the conformation of this region and increased the residue coverage from 67.5% to 90.7% and the lDDT from 0.579 to 0.679.

**Figure 4.**
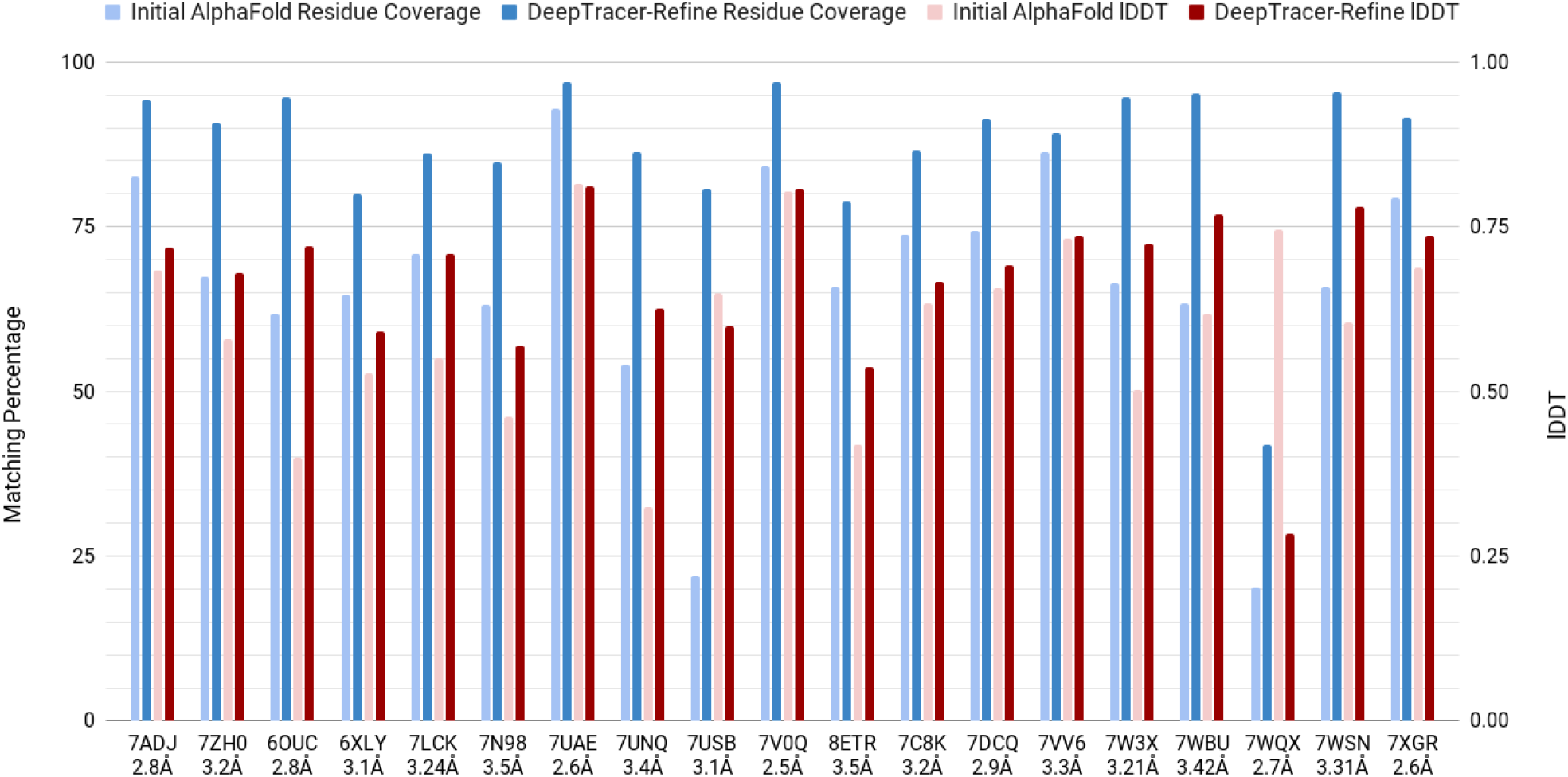
DeepTracer-Refine improvements in residue coverage (left y-axis, 0–100%) and lDDT score (right y-axis, 0.00–1.00). The residue coverage is displayed as blue bars and the lDDT as red bars. The lighter colors represent initial AlphaFold results and the darker colors are DeepTracer-Refine results respectively.

**Figure 5.**
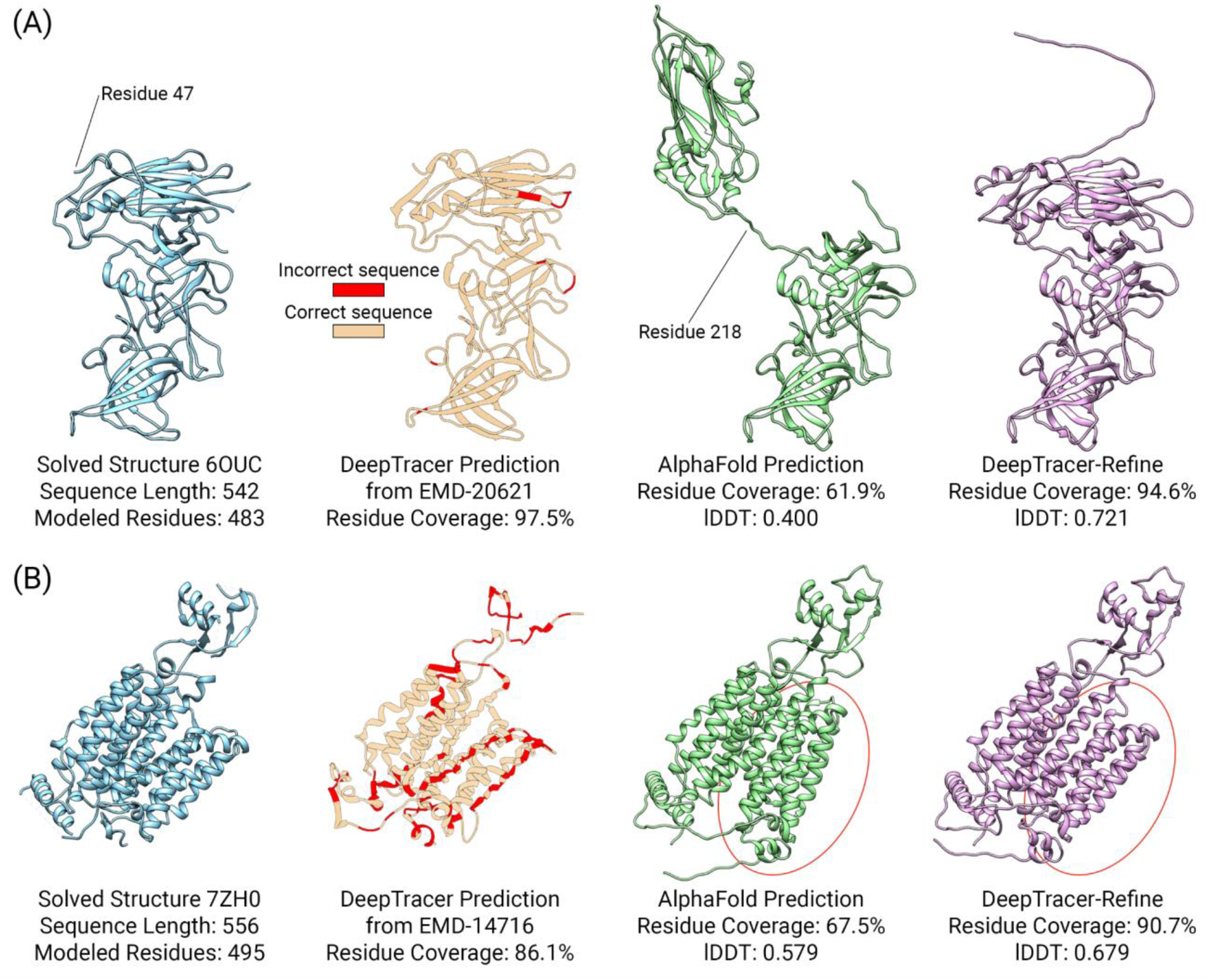
Examples of structural improvements after DeepTracer-Refine. In both examples, the solved and DeepTracer structures do not have complete residue coverage. On the other hand, AlphaFold and DeepTracer-Refine structures contain the entire sequence but the backbones are not accurate enough for the residues to be matched so the residue coverage is incomplete as well. **(A)** The original DeepTracer prediction is accurate in residue coverage and predicted sequence. DeepTracer-Refine split the AlphaFold model at residue 218 and aligned each half correctly. **(B)** An example when the original DeepTracer is not as accurate in sequence prediction. DeepTracer-Refine improved the encircled region to be more similar with the solved structure.

### Comparison against Phenix-Refine

Next, we compared DeepTracer-Refine to a similar method from the PHENIX software suite which we will refer to as Phenix-Refine [34]. It consists of two main parts, which includes docking-and-rebuilding AlphaFold’s prediction from a cryo-EM density map, and utilizing AlphaFold to refold the rebuilt structure. Since Terwilliger et al. have already shown that iterative refolding using the rebuilt structures as AlphaFold templates can further improve the predicted structure, we only compared DeepTracer-Refine to the docking-and-rebuilding portion of their method. The first step is the `Process Predicted Model` method which removes any residues with a pLDDT score of 0.700 or less and groups the remaining residues into three compact domains. The second step is using `Dock Predicted Model` to fit the three compact domains into the cryo-EM density map. Lastly, `Rebuild Predicted Model` traces the trimmed backbone from the density map.

On average, DeepTracer-Refine achieved a residue coverage of 90.0%, which slightly exceeded Phenix-Refine’s 88.4%. On the other hand, Phenix-Refine improved the average lDDT to 0.728 as opposed to DeepTracer-Refine’s 0.707. We attribute the lesser lDDT score to the four structures that worsened after DeepTracer-Refine. This indicates that Phenix-Refine is overall marginally better at improving AlphaFold structures. However, we observe that DeepTracer-Refine is more effective at improving structures with poorer initial results. In Figure 6, we compare the results of both refinement methods to the original AlphaFold predictions and observe that DeepTracer-Refine generally outperforms Phenix-Refine when the initial residue coverage is less than 80% and the initial lDDT is less than 0.65. Furthermore, Phenix-Refine takes magnitudes longer for the docking and rebuilding methods to complete. DeepTracer-Refine outperformed Phenix-Refine in run-time on all testing targets and this is mainly due to DeepTracer utilizing the graphics card to accelerate the backbone tracing process.

**Figure 6.**
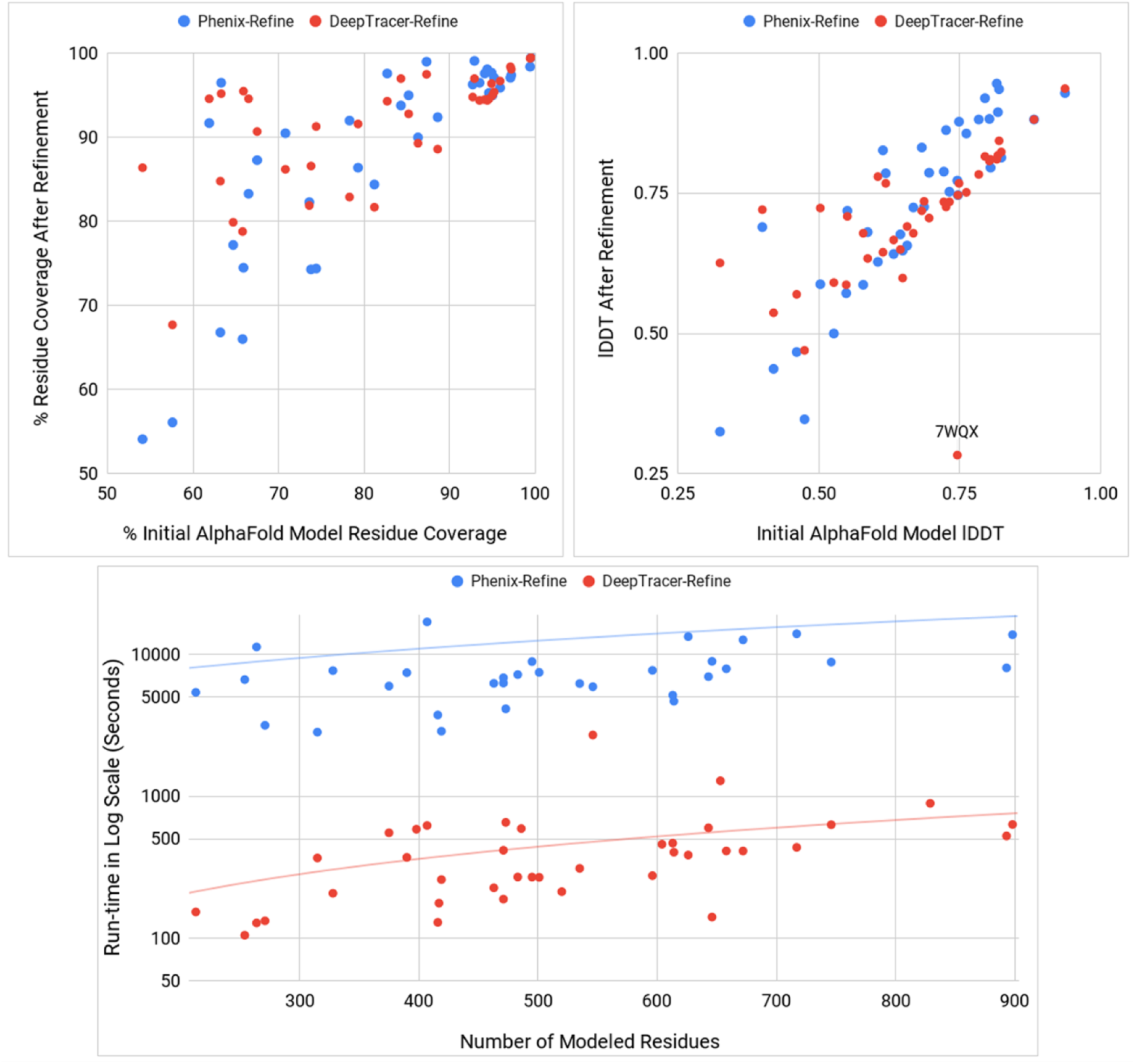
**Top right** and **Top left** graphs compare the results of DeepTracer-Refine and Phenix-Refine to AlphaFold’s initial predicted structures. DeepTracer-Refine is more effective when the initial AlphaFold prediction is less accurate (residue coverage < 80% and lDDT < 0.65). **Bottom** graph compares the run-time performance and it is displayed in logarithmic scale. The average run-time of DeepTracer-Refine is approximately 8 minutes while Phenix-Refine is 3 hours and 35 minutes.

## Discussion

There were four targets that improved in residue coverage after DeepTracer-Refine but decreased in lDDT score, which indicates worsened structures. There are two main reasons why DeepTracer-Refine was unable to improve the initial AlphaFold structure. The first was caused by missing residues in DeepTracer’s prediction, such as 7USB [35] and 7WQX. The corresponding density map EMD-26731 contained weak-density regions so DeepTracer did not trace the backbone properly (Figure 7). This is evident when comparing the solved structure to the AlphaFold prediction. The sequence length is 337 but only 213 residues were modeled in the solved structure. In this instance, DeepTracer-Refine could not correctly align the compact domains due to the absence of backbone geometry. The second type of worsened result occurs when AlphaFold’s initial prediction is not accurate enough. DeepTracer-Refine was designed to address incorrect movements between domains but not designed to fix imprecise folds within a protein domain.

**Figure 7.**
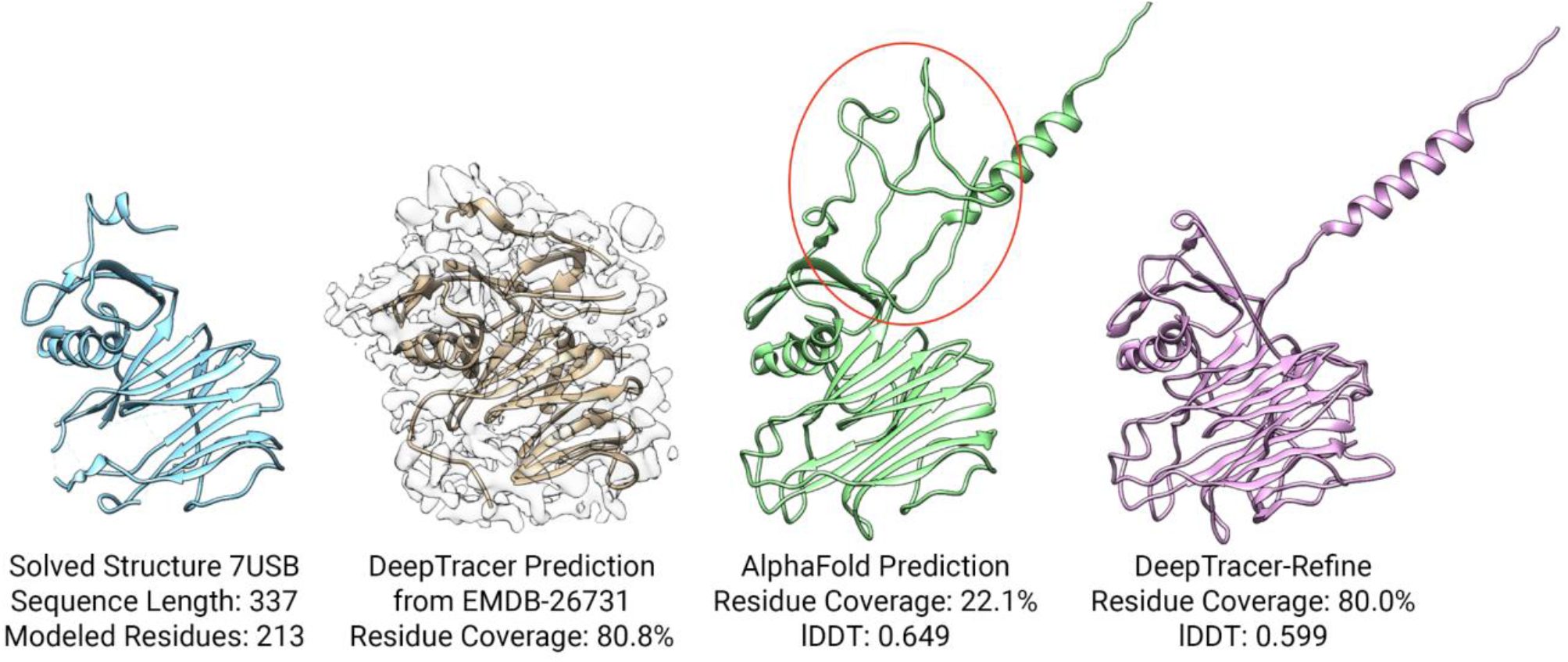
An example of a worsened AlphaFold structure after DeepTracer-Refine. This is caused by the missing backbone in the DeepTracer prediction (highlighted in the red circle) so the compact domains were unable to be properly aligned.

We have shown that the prediction of DeepTracer can be employed to improve the imprecise movements of AlphaFold’s compact domains. However, we did not refine the backbone on an individual residue level due to the challenge in mapping residues between AlphaFold and DeepTracer structures. Both our method and Phenix-Refine only utilize cryo-EM density maps for compact domain docking, and therefore, we are indirectly supplementing structural information to AlphaFold’s predictions. If we are able to accurately match residues from a map-to-model structure onto a sequence, then we could supplement the structural information directly from a cryo-EM density map to AlphaFold. As such, AlphaFold would be able to reference spatial information from experimental density maps to improve the low-confidence folds. In early 2023, Model Angelo has not only addressed the issue of structure-to-sequence residue matching but also provided a method to refine the backbone geometry of a predicted structure [36]. We are inspired by its backbone refinement module and we are working on a method to refine the backbone of a sequence-to-model structure on an individual residue level.

## Conclusion

Currently, protein structure modeling using AlphaFold2 predictions still frequently requires manual adjustments to attain the final result. Meanwhile, map-to-model methods like DeepTracer are quite useful but it does not take full advantage of the true sequence. Therefore, it is a practical application to incorporate the two approaches to address the shortcomings of each other. In this paper, we presented and tested an automated method that improved the overall conformation of AlphaFold’s initial prediction by utilizing DeepTracer. Our main contribution is proposing a method to quantify the poorness of low-confidence locations from an AlphaFold prediction and employ it to reliably detect and rank optimal splitting locations. Our second contribution is demonstrating that even though DeepTracer’s predictions contain faults, they are still very effective for aligning the compact domains using various alignment algorithms. We have shown that it is not necessary to use cryo-EM electron density maps to dock the compact domains and we can speed up the run-time as a result. We tested our method on 39 multi-domain protein structures and compared it to AlphaFold. We increased the average residue coverage from 78.2% to 90.0% and average lDDT score from 0.67 to 0.71, which indicates a significant improvement in structural similarity to the solved structures.

