## Supplemental Tables for "Protein Structure Refinement via DeepTracer and AlphaFold2"

**Supplementary Material**

**Table 1: A list of 39 proteins used in testing DeepTracer-Refine and Phenix-Refine**

| EMDB ID | PDB ID | Resolution | Domains Based on SWORD2 [37] | Sequence Length | Modeled Residues | % DeepTracer Residue Coverage | % DeepTracer Sequence |
|---|---|---|---|---|---|---|---|
| 6239 | 3J9D | 3.3 | 3 | 961 | 746 | 99.2 | 94.37 |
| 6240 | 3J9E | 3.3 | 1 | 526 | 520 | 84.2 | 82.88 |
| 10857 | 6YNV | 2.8 | 2 | 480 | 417 | 99.8 | 98.32 |
| 10940 | 6YV1 | 3.4 | 1 | 487 | 328 | 97.3 | 78.35 |
| 11727 | 7ADJ | 2.8 | 3 | 732 | 613 | 90 | 56.28 |
| 11731 | 7ADM | 3.5 | 3 | 750 | 614 | 83.6 | 38.76 |
| 12604 | 7NVH | 3 | 1 | 761 | 717 | 96.9 | 90.79 |
| 12638 | 7NXF | 3.1 | 4 | 920 | 829 | 81.1 | 59.11 |
| 13581 | 7PPJ | 3.44 | 4 | 919 | 643 | 88.2 | 27.84 |
| 14618 | 7ZC2 | 2.72 | 2 | 485 | 471 | 95.3 | 92.36 |
| 14716 | 7ZH0 | 3.2 | 2 | 556 | 495 | 86.1 | 59.8 |
| 20202 | 6OUC | 2.8 | 3 | 542 | 483 | 97.5 | 93.17 |
| 20621 | 6U2K | 2.93 | 2 | 642 | 604 | 90.9 | 75 |
| 20922 | 6UWF | 3.08 | 2 | 422 | 416 | 96.6 | 94.94 |
| 22014 | 6X2Z | 3.03 | 1 | 526 | 419 | 96.9 | 89.26 |
| 22252 | 6XLY | 3.1 | 3 | 699 | 646 | 91 | 70.43 |
| 23276 | 7LCK | 3.24 | 2 | 491 | 390 | 90.5 | 55.38 |
| 23482 | 7LQ6 | 3.28 | 2 | 747 | 672 | 91.2 | 70.83 |
| 24252 | 7N98 | 3.5 | 2 | 534 | 473 | 92.6 | 69.13 |
| 24731 | 7RXH | 2.3 | 3 | 735 | 626 | 98.2 | 97.12 |
| 25401 | 7SRQ | 2.7 | 1 | 311 | 254 | 91.3 | 55.51 |
| 25648 | 7T32 | 3.4 | 2 | 391 | 375 | 88.3 | 68.8 |
| 26155 | 7TX6 | 3.3 | 2 | 678 | 398 | 90.7 | 70.35 |
| 26422 | 7UAE | 2.6 | 3 | 587 | 535 | 98.7 | 92.15 |
| 26629 | 7UNQ | 3.4 | 4 | 840 | 486 | 91.2 | 65.43 |
| 26731 | 7USB | 3.1 | 1 | 337 | 213 | 80.8 | 37.56 |
| 26948 | 7V0Q | 2.5 | 3 | 691 | 658 | 96.2 | 89.97 |
| 27094 | 8CZC | 2.86 | 2 | 315 | 315 | 66 | 49.21 |
| 28596 | 8ETR | 3.5 | 4 | 543 | 471 | 83 | 49.26 |
| 30306 | 7C8K | 3.2 | 3 | 596 | 596 | 95.6 | 81.71 |
| 30639 | 7DCQ | 2.9 | 3 | 876 | 653 | 93.6 | 74.89 |

| | | | | | | | |
|---|---|---|---|---|---|---|---|
| 32139 | 7VV6 | 3.3 | 1 | 330 | 271 | 94.5 | 87.45 |
| 32295 | 7W3X | 3.21 | 5 | 934 | 893 | 93.4 | 64.84 |
| 32406 | 7WBU | 3.42 | 6 | 932 | 898 | 90.3 | 65.48 |
| 32715 | 7WQX | 2.7 | 3 | 501 | 501 | 41.7 | 6.99 |
| 32761 | 7WSN | 3.31 | 2 | 520 | 463 | 94 | 62.42 |
| 33187 | 7XGR | 2.6 | 5 | 671 | 546 | 92.3 | 74.54 |
| 33555 | 7Y13 | 3.1 | 1 | 446 | 264 | 92 | 80.3 |
| 34176 | 8GOE | 3 | 2 | 544 | 407 | 98.8 | 95.09 |

**Table 2: DeepTracer-Refine vs AlphaFold2 Results**

| EMDB ID | PDB ID | Resolution | % AlphaFold Residue Coverage | % DeepTracer-Refine Residue Coverage | AlphaFold lDDT | DeepTracer-Refine lDDT |
|---|---|---|---|---|---|---|
| 6239 | 3J9D | 3.3 | 94.4 | 94.4 | 0.726 | 0.726 |
| 6240 | 3J9E | 3.3 | 81.2 | 81.7 | 0.645 | 0.65 |
| 10857 | 6YNV | 2.8 | 99.5 | 99.5 | 0.882 | 0.882 |
| 10940 | 6YV1 | 3.4 | 92.7 | 94.8 | 0.722 | 0.735 |
| 11727 | 7ADJ | 2.8 | 82.7 | 94.3 | 0.683 | 0.719 |
| 11731 | 7ADM | 3.5 | 85.2 | 92.8 | 0.668 | 0.679 |
| 12604 | 7NVH | 3 | 87.3 | 97.5 | 0.795 | 0.816 |
| 12638 | 7NXF | 3.1 | 73.6 | 81.9 | 0.549 | 0.587 |
| 13581 | 7PPJ | 3.44 | 97.2 | 98.1 | 0.824 | 0.824 |
| 14618 | 7ZC2 | 2.72 | 94.9 | 96.4 | 0.82 | 0.844 |
| 14716 | 7ZH0 | 3.2 | 67.5 | 90.7 | 0.579 | 0.679 |
| 20202 | 6OUC | 2.8 | 61.9 | 94.6 | 0.4 | 0.721 |
| 20621 | 6U2K | 2.93 | 93.5 | 94.4 | 0.749 | 0.768 |
| 20922 | 6UWF | 3.08 | 57.6 | 67.7 | 0.475 | 0.47 |
| 22014 | 6X2Z | 3.03 | 95.9 | 96.7 | 0.805 | 0.811 |
| 22252 | 6XLY | 3.1 | 64.7 | 79.9 | 0.527 | 0.591 |
| 23276 | 7LCK | 3.24 | 70.8 | 86.2 | 0.551 | 0.709 |
| 23482 | 7LQ6 | 3.28 | 78.3 | 82.9 | 0.587 | 0.634 |
| 24252 | 7N98 | 3.5 | 63.2 | 84.8 | 0.461 | 0.57 |
| 24731 | 7RXH | 2.3 | 95.2 | 95.4 | 0.818 | 0.818 |
| 25401 | 7SRQ | 2.7 | 94.1 | 94.5 | 0.614 | 0.645 |
| 25648 | 7T32 | 3.4 | 97.1 | 98.4 | 0.762 | 0.752 |
| 26155 | 7TX6 | 3.3 | 95 | 95 | 0.747 | 0.747 |

| | | | | | | |
|---|---|---|---|---|---|---|
| 26422 | 7UAE | 2.6 | 92.9 | 97 | 0.816 | 0.811 |
| 26629 | 7UNQ | 3.4 | 54.1 | 86.4 | 0.325 | 0.626 |
| 26731 | 7USB | 3.1 | 22.1 | 80.8 | 0.649 | 0.599 |
| 26948 | 7V0Q | 2.5 | 84.3 | 97 | 0.803 | 0.808 |
| 27094 | 8CZC | 2.86 | 99.4 | 99.4 | 0.937 | 0.937 |
| 28596 | 8ETR | 3.5 | 65.8 | 78.8 | 0.42 | 0.537 |
| 30306 | 7C8K | 3.2 | 73.8 | 86.6 | 0.633 | 0.667 |
| 30639 | 7DCQ | 2.9 | 74.4 | 91.3 | 0.657 | 0.691 |
| 32139 | 7VV6 | 3.3 | 86.3 | 89.3 | 0.732 | 0.735 |
| 32295 | 7W3X | 3.21 | 66.5 | 94.6 | 0.503 | 0.724 |
| 32406 | 7WBU | 3.42 | 63.3 | 95.2 | 0.619 | 0.768 |
| 32715 | 7WQX | 2.7 | 20.2 | 41.9 | 0.746 | 0.283 |
| 32761 | 7WSN | 3.31 | 65.9 | 95.5 | 0.605 | 0.78 |
| 33187 | 7XGR | 2.6 | 79.3 | 91.6 | 0.687 | 0.736 |
| 33555 | 7Y13 | 3.1 | 88.6 | 88.6 | 0.696 | 0.706 |
| 34176 | 8GOE | 3 | 94.6 | 94.6 | 0.784 | 0.784 |

**Table 3: DeepTracer-Refine vs Phenix-Refine Results**

| EMDB ID | PDB ID | % DeepTracer-Refine Residue Coverage | % Phenix-Refine Residue Coverage | DeepTracer-Refine lDDT | Phenix-Refine lDDT | DeepTracer-Refine Run-time (sec) | Phenix-Refine Run-time (sec) |
|---|---|---|---|---|---|---|---|
| 6239 | 3J9D | 94.4 | 98.1 | 0.726 | 0.863 | 631.07 | 8847.88 |
| 6240 | 3J9E | 81.7 | 84.4 | 0.65 | 0.677 | 212.93 | 124131.29 |
| 10857 | 6YNV | 99.5 | 99.5 | 0.882 | 0.882 | 176.55 | N/A |
| 10940 | 6YV1 | 94.8 | 96.3 | 0.735 | 0.789 | 207.34 | 7698.94 |
| 11727 | 7ADJ | 94.3 | 97.6 | 0.719 | 0.832 | 468.29 | 5170.65 |
| 11731 | 7ADM | 92.8 | 95 | 0.679 | 0.725 | 403.81 | 4689.74 |
| 12604 | 7NVH | 97.5 | 99 | 0.816 | 0.92 | 436.61 | 14017.95 |
| 12638 | 7NXF | 81.9 | 82.3 | 0.587 | 0.572 | 893.08 | 24213.08 |
| 13581 | 7PPJ | 98.1 | 97.4 | 0.824 | 0.814 | 599.25 | 6980.46 |
| 14618 | 7ZC2 | 96.4 | 97.7 | 0.844 | 0.936 | 188.53 | 6290.08 |
| 14716 | 7ZH0 | 90.7 | 87.3 | 0.679 | 0.587 | 269.62 | 8920.88 |
| 20202 | 6OUC | 94.6 | 91.7 | 0.721 | 0.69 | 269.96 | 7222.1 |
| 20621 | 6U2K | 94.4 | 96.5 | 0.768 | 0.878 | 459.89 | 57290.51 |
| 20922 | 6UWF | 67.7 | 56.1 | 0.47 | 0.347 | 129.12 | 3749.59 |
| 22014 | 6X2Z | 96.7 | 95.9 | 0.811 | 0.796 | 259 | 2879.38 |
| 22252 | 6XLY | 79.9 | 77.2 | 0.591 | 0.5 | 140.85 | 8955.14 |

| | | | | | | | |
|---|---|---|---|---|---|---|---|
| 23276 | 7LCK | 86.2 | 90.5 | 0.709 | 0.719 | 371.49 | 7443.99 |
| 23482 | 7LQ6 | 82.9 | 92 | 0.634 | 0.681 | 412.11 | 12696.55 |
| 24252 | 7N98 | 84.8 | 66.8 | 0.57 | 0.467 | 655.31 | 4144.53 |
| 24731 | 7RXH | 95.4 | 97.1 | 0.818 | 0.895 | 385.35 | 13376.77 |
| 25401 | 7SRQ | 94.5 | 97.6 | 0.645 | 0.827 | 104.76 | 6653.01 |
| 25648 | 7T32 | 98.4 | 97.1 | 0.752 | 0.857 | 553.25 | 5989.05 |
| 26155 | 7TX6 | 95 | 95 | 0.747 | 0.747 | 586.4 | N/A |
| 26422 | 7UAE | 97 | 99.1 | 0.811 | 0.946 | 310.41 | 6242.24 |
| 26629 | 7UNQ | 86.4 | 54.1 | 0.626 | 0.325 | 592.26 | N/A |
| 26731 | 7USB | 80.8 | 91.1 | 0.599 | 0.648 | 153.11 | 5408.96 |
| 26948 | 7V0Q | 97 | 93.8 | 0.808 | 0.883 | 412.31 | 7938.47 |
| 27094 | 8CZC | 99.4 | 98.4 | 0.937 | 0.929 | 368 | 2835.74 |
| 28596 | 8ETR | 78.8 | 66 | 0.537 | 0.437 | 416.87 | 6856.2 |
| 30306 | 7C8K | 86.6 | 74.3 | 0.667 | 0.642 | 275.78 | 7731.31 |
| 30639 | 7DCQ | 91.3 | 74.4 | 0.691 | 0.657 | 1287.01 | N/A |
| 32139 | 7VV6 | 89.3 | 90 | 0.735 | 0.753 | 132.46 | 3166.41 |
| 32295 | 7W3X | 94.6 | 83.3 | 0.724 | 0.588 | 525.74 | 8056.92 |
| 32406 | 7WBU | 95.2 | 96.5 | 0.768 | 0.786 | 633.81 | 13807.25 |
| 32715 | 7WQX | 41.9 | 94.8 | 0.283 | 0.773 | 268.62 | 7476.64 |
| 32761 | 7WSN | 95.5 | 74.5 | 0.78 | 0.628 | 226.39 | 6262.86 |
| 33187 | 7XGR | 91.6 | 86.4 | 0.736 | 0.726 | 2709.39 | 5929.03 |
| 33555 | 7Y13 | 88.6 | 92.4 | 0.706 | 0.787 | 127.96 | 11318.59 |
| 34176 | 8GOE | 94.6 | 95.3 | 0.784 | 0.882 | 622.5 | 16998.94 |
